# A meta-analysis of gRNA library screens enables an improved understanding of the impact of gRNA folding and structural stability on CRISPR-Cas9 activity

**DOI:** 10.1101/2021.05.29.446220

**Authors:** E.A. Moreb, Michael D. Lynch

## Abstract

CRISPR systems are known to be inhibited by unwanted secondary structures that form within the guide RNA (gRNA). The minimum free energy of predicted secondary structures has been used in prediction algorithms. However, the types of structures as well as the degree to which a predicted structure can inhibit Cas9/gRNA activity is not well characterized. Here we perform a meta-analysis of published CRISPR-Cas9 datasets to better understand the role of secondary structures in inhibiting gRNA activity. We identify two inhibitory structures and provide estimated free energy cutoffs at which they become impactful. Further, we identify the prevalence of these structures in existing datasets. The cutoffs provided help to explain conflicting impacts of free energy values in different datasets as well as providing a guideline for future gRNA designs.

**Highlights:**

- Clearly define two secondary structures that inhibit CRISPR-Cas9 activity
- Provide free energy calculations and cutoffs at which each structure begins to inhibit activity
- Evaluate impact of these structures in published datasets

## Introduction

CRISPR-Cas9 on-target activity is dependent on the sequence of the guide RNA (gRNA).^1–3^The mechanisms underlying this sequence dependence have been studied extensively but many unknowns remain.^3–7^ One factor reported to negatively impact CRISPR-Cas9 on-target activity is the formation of unwanted secondary structures.^7,8^ Unwanted RNA structures and their associated stability can be estimated computationally.^9^ The stability of secondary structures in both the spacer sequence and the full gRNA sequence (including scaffold) have been estimated and have been correlated with activity. This approach is incorporated into some predictive algorithms.^5,7^ However, while gRNA structure is known to inhibit activity, it has not been clear which structures impact activity, to what degree, and how prevalent or impactful this is in routine experimentation. We therefore sought to better define inhibitory gRNA structures and their impact in genome editing studies. To do so we expanded upon a meta-analysis we have recently reported, which includes 39 published datasets with gRNA libraries in various organisms.^6^ We report two types of secondary structure that can inhibit Cas9 activity, provide estimated free energy cutoffs at which they begin to inhibit activity, and evaluate their prevalence within published datasets.

## Methods

Datasets were compiled as described previously.^6^ All calculations and generation of figures were performed in Python, using standard libraries.^10–14^ Structure and minimum free energy predictions were calculated using ViennaRNA RNAfold package.^9^ To calculate the average accessibility of each position along the spacer (Figure 1e), we first calculated the predicted secondary structure of the gRNA and scaffold together (Full Structure). Positions in the spacer region of the Full Structure were then assigned a 1 if predicted to be unbound or a 0 if bound and for each group of gRNA (ie, Functional gRNA), we averaged the values at each position within the gRNA. For example, the accessibility at position 1 within the Functional gRNA is the average accessibility at that site for all ∼1.17 million Functional gRNA in the dataset. All code is provided as a Jupyter Notebook in Supplementary File S1.

**Figure 1:**
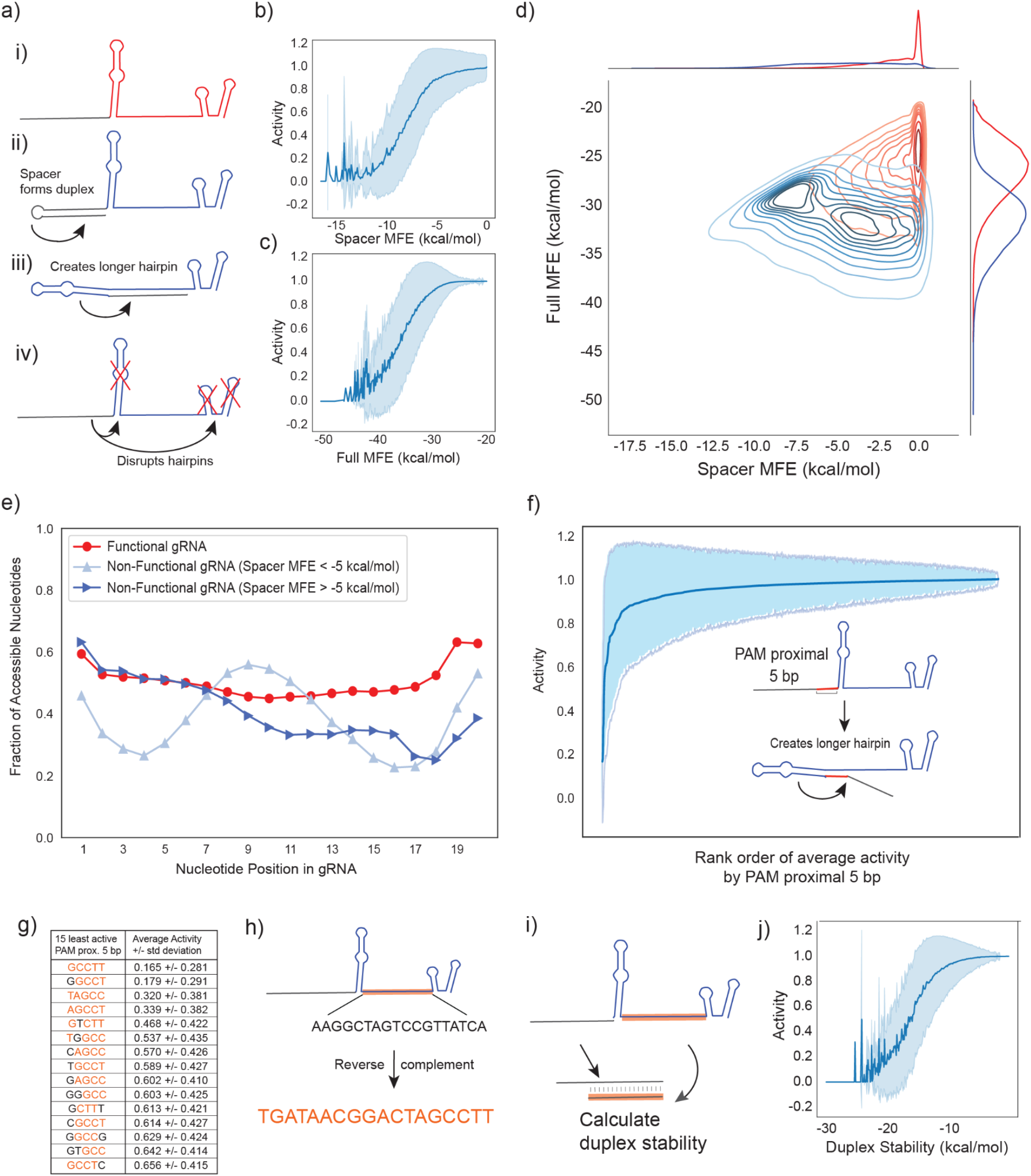
a) Possible inhibitory gRNA structures. Predicted minimum free energy (MFE) of folding of the b) spacer alone (Spacer MFE) and c) the spacer plus scaffold (Full MFE) both show sigmoidal impact on activity. d) Spacer MFE plotted against Full MFE after splitting the dataset into Functional (red) and Non-Functional (blue) gRNA with activity above or below 0.5, respectively. e) Accessibility of each position in the spacer for Functional gRNA (red), Nonfunctional gRNA with Spacer MFE below -5 kcal/mol (light blue), and Nonfunctional gRNA with Spacer MFE above -5 kcal/mol (dark blue). Accessible nucleotides are assigned a 1, bound nucleotides are assigned a 0, and the averaged value for each position is shown. f) All gRNA were grouped by the PAM proximal 5bp of the gRNA and average activity was calculated. Averages are presented by rank order of the PAM proximal 5bp. g) The 15 least active 5bp sequences all have homology to h) the non-structured portion of the gRNA scaffold. i) We calculated Duplex Stability of the spacer sequence binding to the non-structured sequence of the scaffold and k) show a sigmoidal impact on activity. All data presented here is from wild-type Cas9 in the Talas et al dataset.^8^

## Results & Discussion

Of the 39 datasets analyzed, a recent study in *E. coli* provides a unique opportunity to better understand the impact of gRNA structure on Cas9 activity.^8^ Talas et al. 2021 measured the activity of ∼1.2 million self-targeting gRNA (stgRNA) using a plasmid based screening approach in *E. coli*, which enables “the major fraction of the plasmids [to] be cleaved”, resulting in a strongly binary dataset skewed towards active gRNA. Of the ∼1.2 million gRNA, 86.8% had a perfect activity score of 1 while only 4.3% of gRNA are defined as inactive (activity score <0.5). The authors also noted a strong correlation between activity in this screen and the Minimum Free Energy (MFE) predictions of potential secondary structures of both the spacer sequence as well as the full gRNA sequence.^15^ We therefore hypothesized that gRNA that don’t cleave in this study are “defective”. In the event that unwanted hairpins (lower MFE values) inhibit the Cas9-gRNA complex, it makes sense that the activity in this study is highly correlated with MFE values.

Compared to the desired gRNA structure (Figure 1a, structure i), several structures could inhibit activity: hairpins within the spacer (structure ii), binding between the spacer and the non-structured sequence of the scaffold (structure iii), or binding between the spacer and the scaffold such that the natural hairpins of the scaffold are disrupted (structure iv). Structures ii and iii have been reported previously to impact gRNA activity.^7^ To derive inhibitory gRNA structures from these data, we first used the ViennaRNA RNAFold package^9^ to predict the self-folding structures of just the spacer (referred to as Spacer Structure) as well as the full length gRNA (including both the spacer and the scaffold, referred to as Full Structure). We also calculated the MFE for the Spacer Structure (Spacer MFE) and the Full Structure (Full MFE). We saw a strong sigmoidal impact on activity in the dataset from Talas et al., for both the Spacer MFE and Full MFE (Figure 1b and 1c, respectively).^8^ However, the Full MFE also contains the spacer sequence and therefore strong hairpins in the spacer would also impact the Full MFE results. To better separate these two potential effects, we binned the gRNA activities into two groups, Functional gRNA with an activity score >0.5 and Nonfunctional gRNA with activity <0.5, and compared the relationship between Spacer MFE and Full MFE (Figure 1d). In the Nonfunctional gRNA, two unique populations can be identified, confirming that while there is overlap between structures generating low Spacer and Full MFEs, they collectively capture two unique gRNA structures.

We next turned to better computationally define these two distinct groups of gRNA. In contrast to the Full MFE, the Spacer MFE represents only the stability of structure ii. As a result, we first divided the Nonfunctional gRNA (activity <0.5) into two groups with Spacer MFE values above or below -5 kcal/mol. We then looked at the 20bp spacer region of the Full Structure and assigned either a 1 or 0 to each base position, depending on whether the base was predicted to be bound (0, in a stem loop) or free (1) (Figure 1e). For Functional gRNA, all positions were, on average, accessible. In contrast, the two Nonfunctional gRNA groups both showed reduced accessibility (Figure 1e). For gRNA with a Spacer MFE < -5 kcal/mol, we identified a pattern of inaccessible nucleotides separated by a group of accessible nucleotides, which is consistent with structure ii and expected for strongly negative Spacer MFE values. For the other Nonfunctional gRNA, there is a drop in accessibility in the seed region of the gRNA. This is consistent with either structure iii or iv but suggests that a defining feature of these inhibitory structures is obstruction of the seed region. To help delineate between structure iii and iv, we next grouped all gRNA by the PAM proximal 5bp of the spacer, calculated the average activity for each group, and rank ordered the sequences by average activity (Figure 1f). The least active fifteen 5bp sequences all have strong homology to the non-structured sequence of the gRNA scaffold, consistent with structure iii (Figure 1g-h). Notably, this explains a previously reported but unexplained inhibitory GCC motif in the PAM proximal 5bp sequence.^8,16^ Furthermore, we used RNAFold^9^ to calculate the duplex stability between the spacer and non-structured sequence of the scaffold (hereafter referred to as Duplex Stability, Figure 1i). We see a sigmoidal relationship between activity and Duplex Stability (Figure 1j). This both confirms structure iii as inhibitory of Cas9 activity and provides a method for predicting this structure that is not confounded by predicted structures within the spacer.

We next wanted to assess the impact of these structures on gRNA activity in a larger group of datasets. As mentioned, we have previously compiled 39 datasets from various species.^6^ We filtered data from these 39 studies and included only libraries with 10,000 or more gRNA and then compared Spacer MFE and gRNA activity across these datasets, including datasets with different Cas9 variants (Figure 2a-f).^5,17–20^ Despite differences between datasets, there is a clear relationship between Spacer MFE values and activity when the Spacer MFE is below -5 kcal/mol but not when it is greater. We repeated this for Duplex Stability, setting a similar cutoff, in this case at -15 kcal/mol (Figure 2g-l). We noted that in some datasets less stable Duplex Stability values (closer to 0) appeared to negatively impact activity, contrary to expectations. This trend appeared to be species dependent, with human datasets most impacted. To better understand this relationship, we compared the GC content of a gRNA to its Duplex Stability and noted a strong correlation, particularly in the range of low GC (Figure 3a). Extreme GC content has previously been reported to negatively impact Cas9 activity in certain datasets.^1,3^ In the human data set from Wang et al 2019^5^, if we remove gRNA with GC content less than 30%, we see improved on-target activity within the same range of less stable Duplex Stability values (Figure 3b). Interestingly, the observation that low GC content reduces gRNA activity is not shared among different species and therefore does not impact the relationship between Duplex Stability and activity in the same manner (Supplemental Figure S1). This result, suggests that this observation is due to other context-dependent factors impacted by sequence composition rather than Duplex Stability.^6^ This supports a model in which Duplex Stability below the cutoff we established is representative of formation of structure iii, while Duplex Stability above the cutoff does not indicate formation of inhibitory structures but rather may be correlated with other sequence-dependent features (Figure 3c).

**Figure 2:**
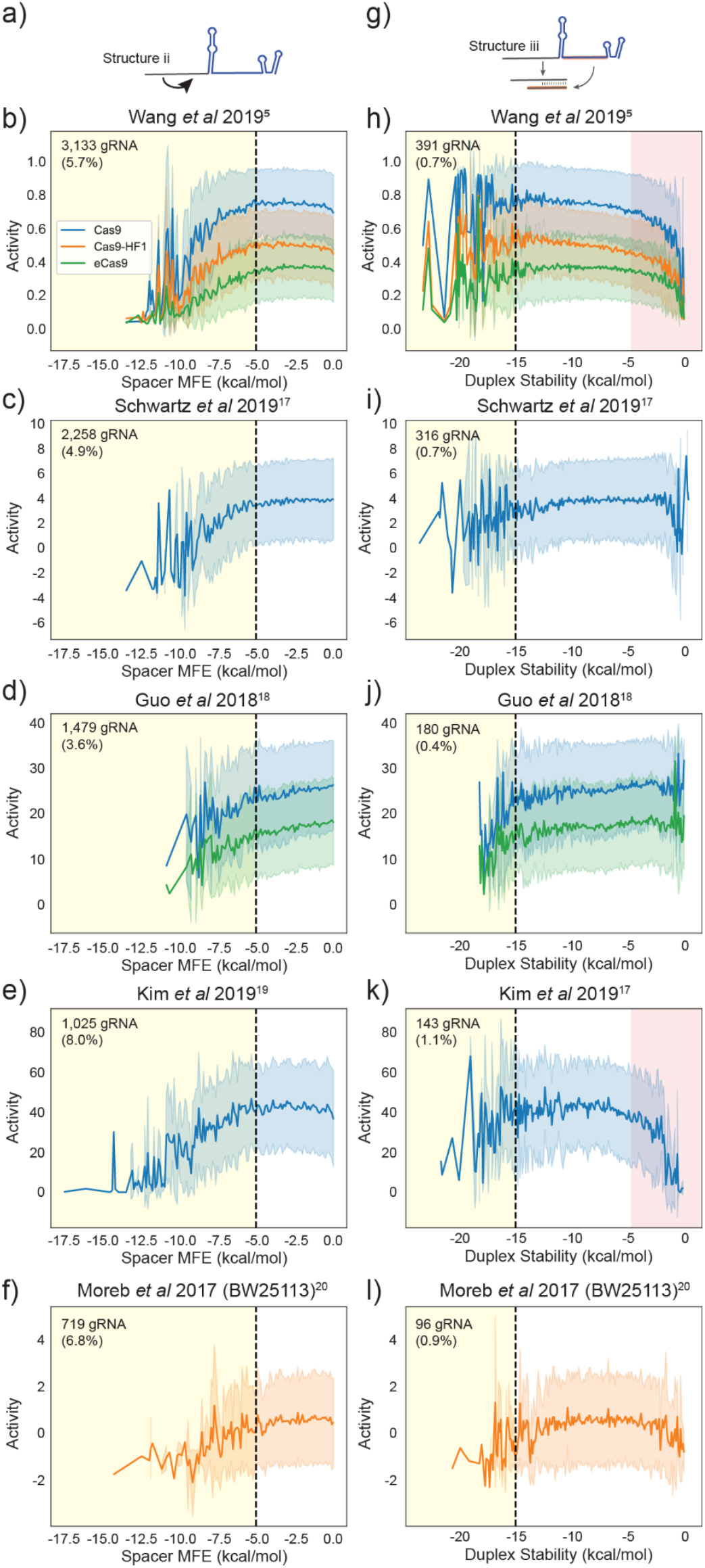
The impact of a) structure ii on gRNA activity is seen in the relationship between Spacer MFE and gRNA activity is highlighted in the five largest datasets with over 10,000 gRNA each (b-f) and across different Cas9 variants (wild-type Cas9: blue, Cas9-HF1: orange, eCas9: green). The yellow shaded region shows gRNA with a Spacer MFE below -5 kcal/mol. Similarly, the impact of g) structure iii, calculated as Duplex Stability, is shown for the same five datasets (h-l) and Cas9 variants. The yellow shaded region highlights gRNA below -15 kcal/mol. The red shaded regions in human datasets highlight an observed negative impact on activity for predicted unstable (low GC or high AT) duplexes, which is contrary to expectations based on structure ii) and iii) alone (see Figure 3).

**Figure 3:**
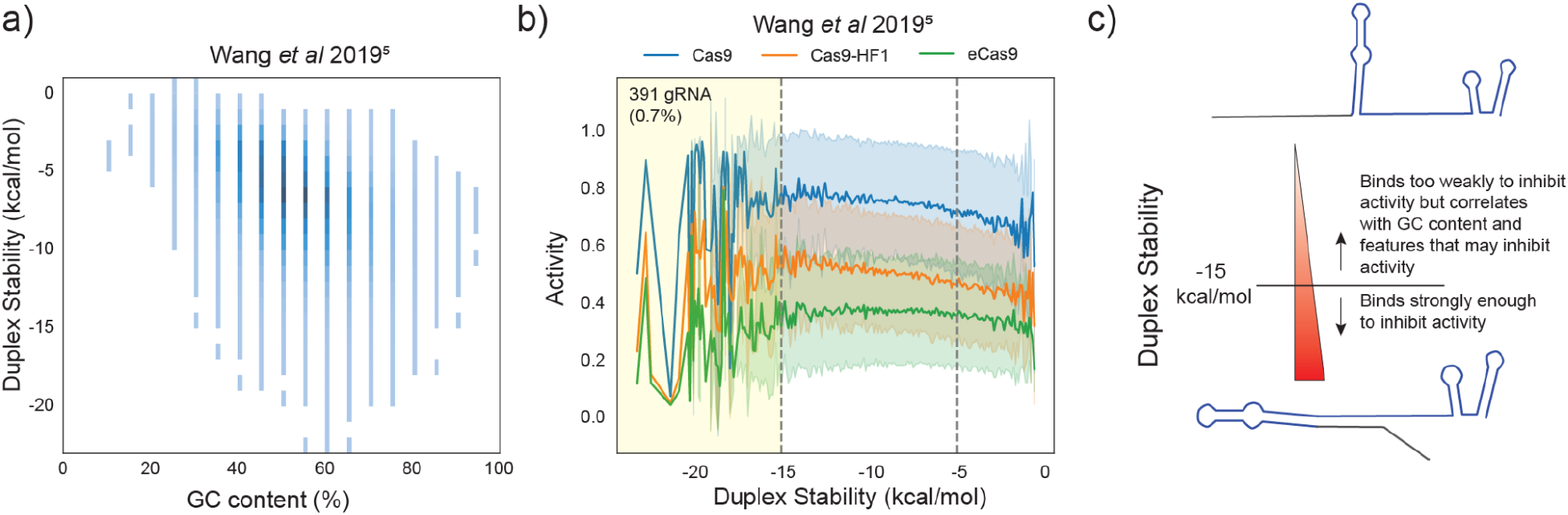
a) Duplex Stability is correlated with gRNA GC content. b) To demonstrate the impact of this correlation on the high range of Duplex Stability, data from Wang et al 2019^5^ were re-plotted without gRNA containing GC content below 30%. c) This supports a model in which Duplex Stability below the cutoff we established is inhibitory based on formation of inhibitory structures, while Duplex Stability above the cutoff does not indicate formation of inhibitory structures but may correlate with other sequence-dependent influences on activity.

Finally, we turned to evaluate the impact of these structures across the remaining smaller datasets. We first calculated the percent of gRNA that are below our proposed cutoffs in each dataset (Figure 4a).^3,4,17,18,20–31^ Overall, structure ii is much more prevalent than structure iii, ranging from 2.7-23.4% of gRNA compared to 0-1.6%, respectively. This variability in relative prevalence of these structures within datasets may have confounding effects on activity predictions. For example, one of the datasets, reported in Doench et al 2014^3^ and used to train the WU-CRISPR algorithm^7^, has 23.4% of gRNA below our Spacer MFE cutoff. In contrast, the second highest percentage is 12.6%. This may lead the WU-CRISPR algorithm to more heavily weight the Spacer MFE. Surprisingly, we still see a large number of datasets published after these studies with a relatively high percent of gRNA predicted to contain structure ii or iii, indicating that a more clear consensus on the impact of structure on activity is needed.

**Figure 4:**
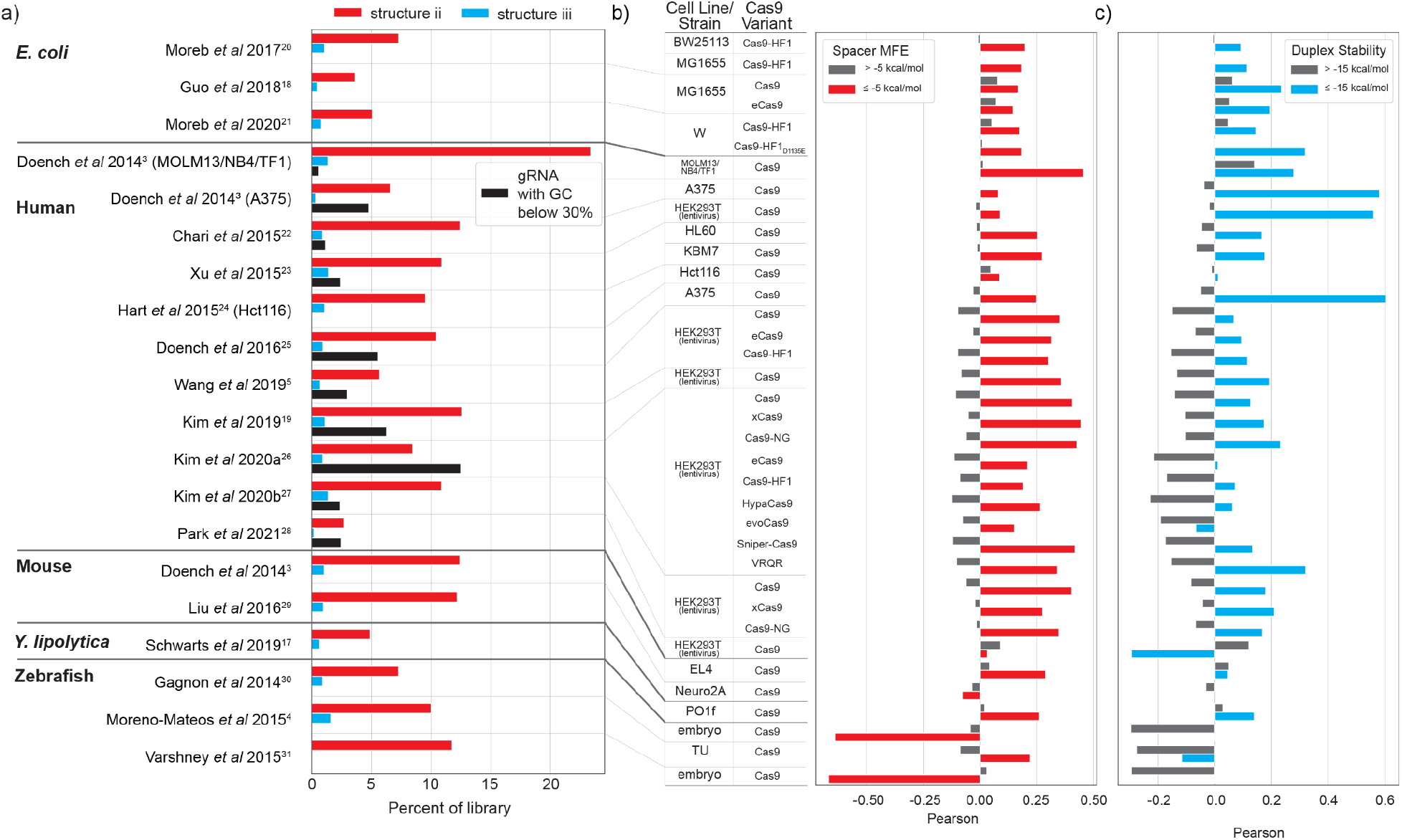
The impact of gRNA structure is highlighted in different datasets. a) The percent of gRNA in each library that contain a predicted structure ii or iii below the relevant cutoffs. For human datasets, we also calculate the percent of gRNA with less than 30% GC. b) Correlation between gRNA activity and Spacer MFE above (grey) and below (red) the -5 kcal/mol cutoff. c) Correlation between gRNA activity and Duplex Stability above (grey) and below (blue) the -15 kcal/mol cutoff.

We next calculated Pearson correlations between Spacer MFE above and below the cutoffs and gRNA activity (Figure 4b). We see broad agreement across datasets that activity is correlated with Spacer MFE below -5 kcal/mol while the correlations above -5 kcal/mol are either reduced or even negative, further highlighting that gRNA with an MFE above -5 kcal/mol are not likely forming structures that inhibit activity. In this regime, other factors are likely contributing to sequence dependent activity. Three datasets show negative correlations with Spacer MFE below - 5 kcal/mol but all three of these datasets have fewer than 205 gRNA and at most 25 gRNA with Spacer MFE below the cutoff. These types of correlations on gRNA libraries this small are unlikely to capture meaningful features. We also calculated Pearson correlations between the Duplex Stability above and below -15 kcal/mol and gRNA activity (Figure 4c). Again, we see several datasets with negative or no correlation between Duplex Stability below -15 kcal/mol and activity. These datasets have relatively few gRNA predicted to have structure iii or are datasets where activity is low overall for other reasons (such as the evoCas9 data from Kim et al 2020a).^26^ Taken together, these results strongly support the use of these new proposed cutoffs for eliminating improperly folded gRNAs in future designs.

In this analysis, we have confirmed two types of secondary structure that can inhibit on-target activity. While previous algorithms for gRNA design have relied on weighted MFE values, this analysis more strongly supports the use of free energy cutoffs to identify “improperly functioning” gRNA. Interestingly, this analysis did not identify structures wherein the natural stem loops of the gRNA scaffold are disrupted. Free energy values above our proposed cutoffs are not meaningfully connected to inhibitory structures but rather may be correlated with other sequence-dependent factors. Furthermore, we have confirmed these cutoffs are practically meaningful across a variety of contexts. These results provide a clear cutoff that can be immediately deployed in new gRNA designs as well as help towards improving algorithms to predict gRNA activity.

## Acknowledgements

We would like to acknowledge the following support: ONR YIP #12043956, and DOE EERE grant #EE0007563. We would also like to acknowledge financial support from Duke Innovation & Entrepreneurship Initiative.

## Author contributions

E.A. Moreb performed computational analyses. E.A. Moreb and M.D. Lynch designed analyses, analyzed results, wrote, revised and edited the manuscript.

## Figures & Captions

**Graphical Abstract**

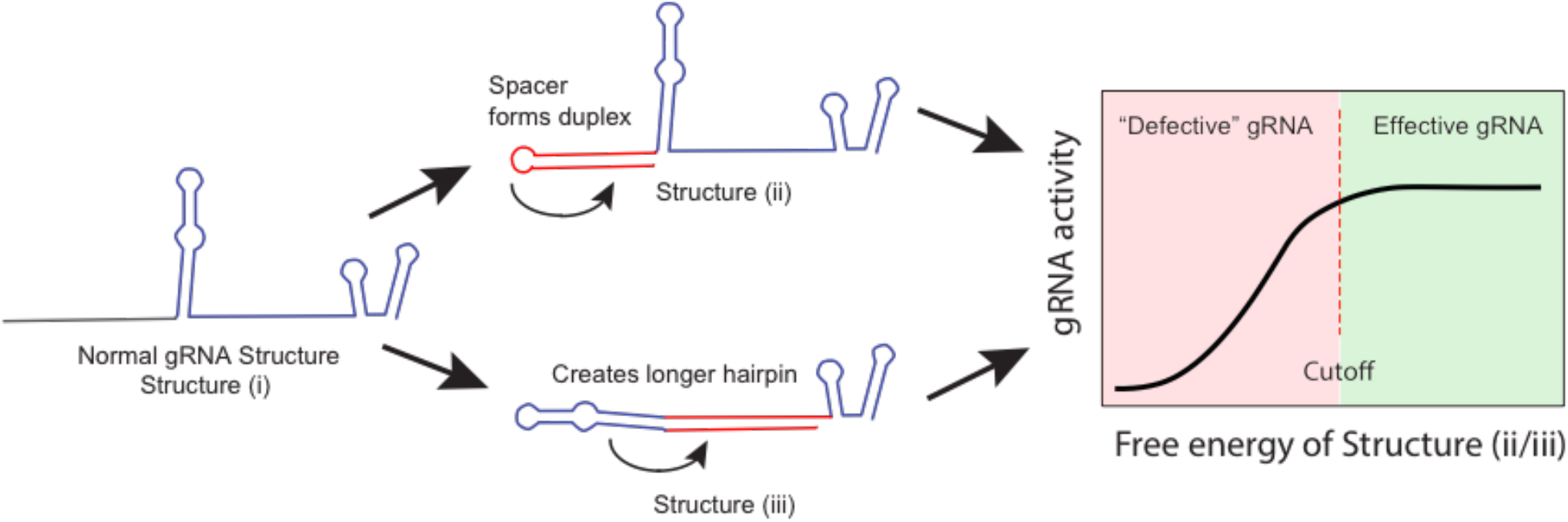

