## Supplementary Materials for "A meta-analysis of gRNA library screens enables an improved understanding of the impact of gRNA folding and structural stability on CRISPR-Cas9 activity"

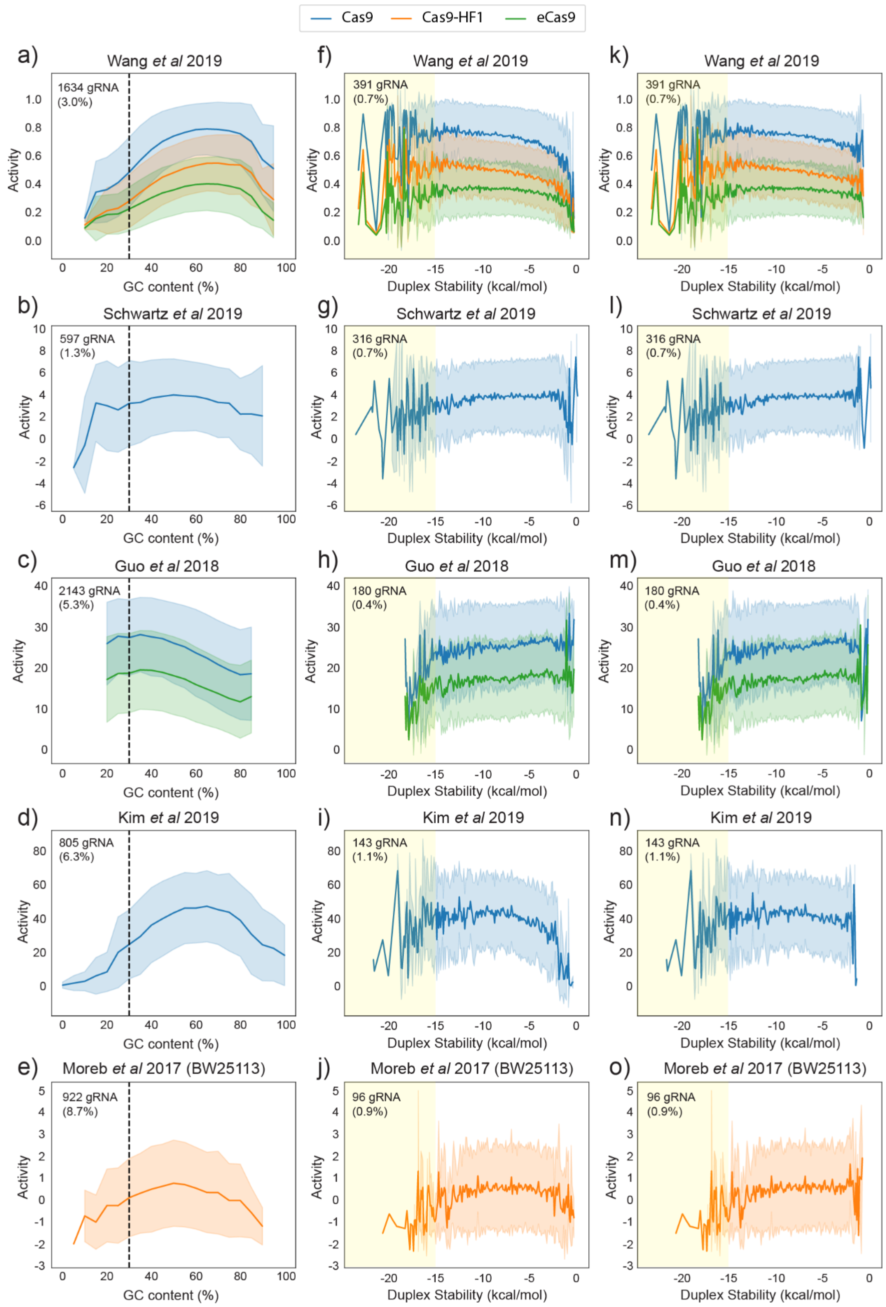


**Supplemental Figure S1:** The relationship between activity and GC content for the five largest datasets is shown in (a-e).Duplex Stability is plotted against activity for the same datasets in (f-j). We removed gRNA with GC below 30% and re-plotted Duplex Stability against activity (k-o).
