## Supplementary File S1 for "A meta-analysis of gRNA library screens enables an improved understanding of the impact of gRNA folding and structural stability on CRISPR-Cas9 activity": Supplemental_File_S1.ipynb

```
# %% [markdown]
**A meta-analysis of gRNA library screens enables an improved understanding of the impact of gRNA folding and structural stability on CRISPR-Cas9 activity.** <br>

Moreb, E.A.*1* , and Lynch, Michael D.*1,2,3* <br>
*1* Department of Biomedical Engineering, Duke University <br>
*2* To whom all correspondence should be addressed. <br>
*3* <br>


# %% [markdown]
**Supplemental Code**

# %% [code]
# Code by E.A. Moreb
#
# Python version 3.8

# %% [markdown]
# Imports/functions

# %% [code]
import numpy as np
import pandas as pd
import matplotlib.pyplot as plt
import seaborn as sns

from matplotlib.collections import LineCollection
from scipy.stats import pearsonr, spearmanr, describe, mannwhitneyu, ttest_ind

from collections import Counter
from itertools import product, repeat
import itertools
from Bio.SeqUtils import MeltingTemp as mt
from Bio.SeqUtils import GC
import re
import math
import RNA # only works on linux or virtual machines

Filepath = # insert filepath

sns.set_style('white')
plt.rcParams["font.family"] = "Arial"

# %% [markdown]
When loading datasets, you will need to rename them.

# %% [code]
tab = str.maketrans("ACTGN","TGACN")

def rev_comp(seq):
    """ Returns the reverse complement of any seuqence. Returns N for N. """
    return seq.translate(tab)[::-1]

dinucleotideOrder = ['TT','AT','CT','GT',
                    'TA','AA','CA','GA',
                    'TC','AC','CC','GC',
                    'TG','AG','CG','GG']

# %% [markdown]
# Loading datasets

# %% [markdown]
These are large files and take a while to load. Datasets from supplemental of Moreb et al 2021

# %% [code]
HumanLibs = pd.read_excel(f"{Filepath}/SupplementalFile2_HumanDatasets.xlsx", 
                        engine='openpyxl')

MouseLibs = pd.read_excel(f"{Filepath}/SupplementalFile4_MouseDatasets.xlsx", 
                        engine='openpyxl')

ColiLibs = pd.read_excel(f"{Filepath}/SupplementalFile3_EcoliDatasets.xlsx", 
                        engine='openpyxl')

ZebraLibs = pd.read_excel(f"{Filepath}/SupplementalFile6_ZebrafishDatasets.xlsx", 
                        engine='openpyxl')

LipolyticaLibs = pd.read_excel(f"{Filepath}/SupplementalFile5_YlipolyticaDatasets.xlsx", 
                              engine='openpyxl')

mergedLibs = pd.concat([HumanLibs, MouseLibs, ColiLibs, ZebraLibs, LipolyticaLibs])

mergedLibs['Dataset_Cas9'] = [f"{x}-{y}" for x,y in mergedLibs[['Dataset','Cas9 Variant']].values.tolist()]
mergedLibs['Year'] = [re.search(r"[0-9]{4,7}", x).group(0) for x in mergedLibs['Dataset'].tolist()]
mergedLibs['30bpSeq'] = [x.upper() for x in mergedLibs['30bpSeq'].tolist()]
print(mergedLibs.shape)
mergedLibs = mergedLibs[mergedLibs['30bpSeq']!='NF'].copy(deep=True) # gRNA Not Found in host genome. Exclude it.
print(mergedLibs.shape)


mergedLibs['NormalizedActivity'] = np.empty(mergedLibs.shape[0])

for ds in mergedLibs.Dataset.unique():
    x = mergedLibs.loc[mergedLibs['Dataset']==ds, 'Activity'].values
    scaler = preprocessing.MinMaxScaler()
    x_scaled = scaler.fit_transform(x.reshape(-1,1))
    mergedLibs.loc[mergedLibs['Dataset']==ds, 'NormalizedActivity'] = x_scaled

# %% [code]
mergedLibs['GC_0:20'] = [GC(x) for x in mergedLibs['gRNA'].tolist()]

# %% [markdown]
Dataset take from supplementary materials of Talas et al 2021

# %% [code]
MillionLib = pd.read_csv(f"{Filepath}/Source Data Fig2 - 1M-library.csv", index_col=0)
MillionLib['WT_binned'] = ['Functional' if x > 0.5 else 'Non-Functional' for x in MillionLib['CutEff_WT'].tolist()]

# %% [code]
MillionLib.head()

# %% [markdown]
# RNAfold calculations

# %% [code]
# mergedLibs MFE_gRNA
mergedLibs['MFE_gRNA']=[RNA.fold(seq)[1] for seq in mergedLibs['gRNA_asExpressed'].tolist()]

# %% [code]
# mergedLibs MFE_gRNAwScaff
mergedLibs['MFE_gRNAwScaff'] = np.empty(mergedLibs.shape[0])
mergedLibs['SS_gRNAwScaff'] = np.empty(mergedLibs.shape[0])

mergedLibs[['SS_gRNAwScaff','MFE_gRNAwScaff']]=[RNA.fold(seq+scaffold) for seq, scaffold in mergedLibs[['gRNA_asExpressed','gRNA Scaffold']].values.tolist()]

# %% [code]
# mergedLibs Duplex Stability
linearScaffold = 'AAGGCTAGTCCGTTATCA' # non-structured portion of the scaffold

mergedLibs['Duplex Stability']=[RNA.duplexfold(seq, linearScaffold).energy for seq in mergedLibs['gRNA_asExpressed'].tolist()]

# %% [code]
# MillionLib MFE_gRNA
MillionLib['MFE_gRNA']=[RNA.fold(seq)[1] for seq in MillionLib['Spacer'].tolist()]

# %% [code]
# MillionLib MFE_gRNAwScaff
selfTargScaffold = "GGGTTAGAGCTAGAAATAGCAAGTTAACCTAAGGCTAGTCCGTTATCAACTTGAAAAAGTGGCACCGAGTCGGTGC"

MillionLib['MFE_gRNAwScaff'] = np.empty(MillionLib.shape[0])
MillionLib['SS_gRNAwScaff'] = np.empty(MillionLib.shape[0])

MillionLib[['SS_gRNAwScaff','MFE_gRNAwScaff']] = [RNA.fold(seq+selfTargScaffold) for seq in MillionLib['Spacer'].tolist()]

# %% [code]
# MillionLib Duplex Stability
MillionLib['Duplex Stability']=[RNA.duplexfold(seq, linearScaffold).energy for seq in MillionLib['Spacer'].tolist()]

# %% [markdown]
# Figure 1

# %% [code]
print('GTTTTAGAGCTAGAAATAGCAAGTTAAAATAAGGCTAGTCCGTTATCAACTTGAAAAAGTGGCACCGAGTCGGTGC') # traditional scaffold
print('(((((((.((((....))))...)))))))..................((((....)))).((((((...))))))') # predicted structure

# %% [markdown]
## Accessibility

# %% [code]
scaffold_colNames = [f"SS_p{i}" for i in np.arange(1,97)]

def plot_fractionAccessible(data, labels, markers=None, colors=None, ax=None):
    """ Data, Labels and Colors are passed as a List to support as many lines as needed. """
    if colors==None:
        colors= ('r','grey','blue')
    if markers == None:
        markers= ('o','^','>')
    if ax:
        for i,ds in enumerate(data):
            ax.plot(ds, label=labels[i], color=colors[i], marker=markers[i])
#         ax.plot(x1, label="Functional sgRNA", marker='o', c=colors[0])
#         ax.plot(x2, label="Non-Functional sgRNA", marker='^', c=colors[1])

        positions = (0, 2, 4, 6, 8, 10, 12, 14, 16, 18)
        labels = ("1", "3", "5", "7", "9", "11", "13", "15", "17", "19")
        ax.set_xticks(positions, labels)

        ax.set_ylim(0,1)
        ax.set_ylabel('Fraction of Accessible Nucleotides')
        ax.set_xlabel('Nucleotide Position in gRNA')
        ax.legend()
    else:
        for i,ds in enumerate(data):
            plt.plot(ds, label=labels[i], color=colors[i], marker=markers[i])
            
#         plt.plot(x1, label="Functional sgRNA", marker='o', c=colors[0])
#         plt.plot(x2, label="Non-Functional sgRNA", marker='^', c=colors[1])

        positions = (0, 2, 4, 6, 8, 10, 12, 14, 16, 18)
        labels = ("1", "3", "5", "7", "9", "11", "13", "15", "17", "19")
        plt.xticks(positions, labels)

        plt.ylim(0,1)
        plt.ylabel('Fraction of Accessible Nucleotides')
        plt.xlabel('Nucleotide Position in gRNA')
        plt.legend()

# %% [code]
for col in scaffold_colNames:
    MillionLib[col] = np.empty(MillionLib.shape[0])
    
MillionLib.loc[:, scaffold_colNames] = pd.DataFrame(index=MillionLib.index, columns=scaffold_colNames, data=[list(x.replace("(","0").replace(")","0").replace(".","1")) for x in MillionLib['SS_gRNAwScaff'].tolist()])

for col in scaffold_colNames:
    MillionLib[col] = pd.to_numeric(MillionLib[col], 'coerce')

# %% [code]
group1 = MillionLib[(MillionLib['CutEff_WT'] >= 0.5) &
                   (MillionLib['MFE_gRNA'] > -5)]
group2 = MillionLib[(MillionLib['CutEff_WT'] < 0.5) &
                   (MillionLib['MFE_gRNA'] > -5)]
group3 = MillionLib[(MillionLib['CutEff_WT'] < 0.5) &
                   (MillionLib['MFE_gRNA'] <= -5)]

x1 = group1.loc[:, scaffold_colNames].mean().tolist()[0:20]
x2 = group2.loc[:, scaffold_colNames].mean().tolist()[0:20]
x3 = group3.loc[:, scaffold_colNames].mean().tolist()[0:20]

plot_fractionAccessible([x1,x2,x3], labels=['Functional gRNA (20bp MFE > -5)',
                                          'Non-Functional gRNA (20bp MFE > -5)',
                                          "Non-Functional sgRNA (20bp MFE < -5)"])


# %% [markdown]
## Sigmoidal plots

# %% [code]
f, axes = plt.subplots(3,1, figsize=(4,12), sharey=True)
sns.lineplot(x='MFE_gRNA', y='CutEff_WT', ci='sd', data=MillionLib, ax=axes[0])
sns.lineplot(x='MFE_gRNAwScaff', y='CutEff_WT', ci='sd', data=MillionLib, ax=axes[1])
sns.lineplot(x='Duplex Stability', y='CutEff_WT', ci='sd', data=MillionLib, ax=axes[2])

# %% [markdown]
## kde plot

# %% [code]
df = MillionLib.copy(deep=True)

Func = df.loc[df.WT_binned=="Functional"]
NonFunc = df.loc[df.WT_binned=="Non-Functional"]

g = sns.JointGrid(x="MFE_gRNA", y="MFE_gRNAwScaff", data=df)
sns.kdeplot(x = "MFE_gRNA", y='MFE_gRNAwScaff', data=Func, cmap="Reds_d", 
            thresh=0.05, levels=10, gridsize=100,
            shade=False,  ax=g.ax_joint)
sns.kdeplot(x = "MFE_gRNA", y='MFE_gRNAwScaff', data=NonFunc, cmap="Blues_d", 
            thresh=0.05, levels=10, gridsize=100,
            shade=False,  ax=g.ax_joint)

sns.kdeplot(x="MFE_gRNA", data=Func, color="r", ax=g.ax_marg_x)
sns.kdeplot(x="MFE_gRNA", data=NonFunc, color="b", ax=g.ax_marg_x)

sns.kdeplot(y="MFE_gRNAwScaff", data=Func, color="r", ax=g.ax_marg_y)
sns.kdeplot(y="MFE_gRNAwScaff", data=NonFunc, color="b", ax=g.ax_marg_y)


# %% [markdown]
## PAM proximal rank order 


# %% [code]
MillionLib['PAMprox5bp'] = [x[-5::] for x in MillionLib['Spacer'].tolist()]

mean_ = MillionLib.groupby('PAMprox5bp')['CutEff_WT'].mean().sort_values()
idx = mean_.index
std_ = MillionLib.groupby('PAMprox5bp')['CutEff_WT'].std().loc[idx]


MillionLib['orderedPAMprox5bp'] = pd.Categorical(MillionLib['PAMprox5bp'], 
                                                categories=idx,
                                                ordered=True)
activityCutoff = 0.8
sns.lineplot(x='orderedPAMprox5bp', y='CutEff_WT', ci='sd', data=MillionLib)


# %% [markdown]
# Figure 2

# %% [code]
# five largest datasets
datasetList = ['Wang2019-293T', 'Schwartz2019-PO1f','Guo2018-MG1655',
               'Kim2019-SA-lenti293T','Moreb2017-BW25113']  


f, axes = plt.subplots(5,2, figsize=(10,20), sharex='col', sharey='row')

cutoff = -5
duplexCutoff = -15

for i, ds in enumerate(datasetList):
    sns.lineplot(x='MFE_gRNA', y='Activity', hue='Cas9 Variant', hue_order=['Cas9','Cas9-HF1','eCas9'],
                 data=mergedLibs[mergedLibs['Dataset']==ds], ci='sd', ax=axes[i,0])
    axes[i,0].set_title(ds)
    axes[i,0].legend(loc='lower left')
    axes[i,0].axvline(cutoff, ls='--', color='k')
    if i < 4 or i > 4:
        dsCas = ds+'-Cas9'
        count = mergedLibs[(mergedLibs['Dataset_Cas9']==dsCas) & (mergedLibs['MFE_gRNA']<=cutoff)].shape[0]
        totalcount = mergedLibs[(mergedLibs['Dataset_Cas9']==dsCas)].shape[0]
        percent= (count/totalcount)*100
        axes[i,0].text(0.05, 0.85, f"{count} gRNA out of \n{totalcount} ({percent: .1f}%) less than {cutoff}",
                    horizontalalignment='left', verticalalignment='center', transform=axes[i,0].transAxes)
    else:
        dsCas = ds+'-Cas9-HF1'
        count = mergedLibs[(mergedLibs['Dataset_Cas9']==dsCas) & (mergedLibs['MFE_gRNA']<=cutoff)].shape[0]
        totalcount = mergedLibs[(mergedLibs['Dataset_Cas9']==dsCas)].shape[0]
        percent= (count/totalcount)*100
        axes[i,0].text(0.05, 0.85, f"{count} gRNA out of \n{totalcount} ({percent: .1f}%) less than {cutoff}",
                    horizontalalignment='left', verticalalignment='center', transform=axes[i,0].transAxes)
    
    
    
    sns.lineplot(x='Duplex Stability', y='Activity', hue='Cas9 Variant', hue_order=['Cas9','Cas9-HF1','eCas9'],
                 data=mergedLibs[mergedLibs['Dataset']==ds], ci='sd', ax=axes[i,1])
    axes[i,1].set_title(ds)
    axes[i,1].legend(loc='lower left')
    axes[i,1].axvline(duplexCutoff, ls='--', c='k')
    
    if i < 4 or i > 4:
        dsCas = ds+'-Cas9'
        count = mergedLibs[(mergedLibs['Dataset_Cas9']==dsCas) & (mergedLibs['Duplex Stability']<=duplexCutoff)].shape[0]
        totalcount = mergedLibs[(mergedLibs['Dataset_Cas9']==dsCas)].shape[0]
        percent= (count/totalcount)*100
        axes[i,1].text(0.05, 0.85, f"{count} gRNA out of \n{totalcount} ({percent: .1f}%) less than {duplexCutoff}",
                    horizontalalignment='left', verticalalignment='center', transform=axes[i,1].transAxes)
    else:
        dsCas = ds+'-Cas9-HF1'
        count = mergedLibs[(mergedLibs['Dataset_Cas9']==dsCas) & (mergedLibs['Duplex Stability']<=duplexCutoff)].shape[0]
        totalcount = mergedLibs[(mergedLibs['Dataset_Cas9']==dsCas)].shape[0]
        percent= (count/totalcount)*100
        axes[i,1].text(0.05, 0.85, f"{count} gRNA out of \n{totalcount} ({percent: .1f}%) less than {duplexCutoff}",
                    horizontalalignment='left', verticalalignment='center', transform=axes[i,1].transAxes)
            


# %% [markdown]
# Figure 3

# %% [code]
ds = 'Wang2019-293T'

f, axes = plt.subplots(1,2, figsize=(10,4))
GCcutoff = 30

sns.histplot(x='GC_0:20', y='Duplex Stability', 
            ax=axes[0], binwidth=(1, 1),
            data=mergedLibs[mergedLibs['Dataset']==ds])
axes[0].set_xlim(0,100)
axes[0].axhline(-5, ls='--', c='grey')
axes[0].axvline(30, ls='--', c='grey')

sns.lineplot(x='Duplex Stability', y='Activity', hue='Cas9 Variant', hue_order=['Cas9','Cas9-HF1','eCas9'],
                 data=mergedLibs[(mergedLibs['Dataset']==ds) & (mergedLibs['GC_0:20']>GCcutoff)], ci='sd', ax=axes[1])


axes[1].axvline(-15, ls='--', c='grey')
axes[1].axvline(-5, ls='--', c='grey')


# %% [markdown]
# Figure 4

# %% [code]
cutoff = -5
GCcutoff = 30
duplexCutoff = -15


test = pd.DataFrame(columns=['Dataset','Type','Percent'])
for ds in mergedLibs[mergedLibs['Dataset']!='Tálas2021-NEB5-alpha'].sort_values(['Genome','Year']).Dataset.unique():
    total = mergedLibs[mergedLibs['Dataset']==ds].shape[0]
    mfe20 = mergedLibs[(mergedLibs['Dataset']==ds) &
                      (mergedLibs['MFE_gRNA']<=cutoff)].shape[0]
    duplexE = mergedLibs[(mergedLibs['Dataset']==ds) &
                      (mergedLibs['Duplex Stability']<=duplexCutoff)].shape[0]
    GC30 = mergedLibs[(mergedLibs['Dataset']==ds) &
                      (mergedLibs['GC_0:20']<=GCcutoff)].shape[0]
    percentmfe20 = (mfe20/total)*100
    percentGC30 = (GC30/total)*100
    percentDupeE = (duplexE/total)*100
    test.loc[test.shape[0]] = [ds, 'struct_ii', percentmfe20]
    test.loc[test.shape[0]] = [ds, 'GC30', percentGC30]
    test.loc[test.shape[0]] = [ds, 'struct_iii', percentDupeE]
    
plt.figure(figsize=(4,10))
sns.barplot(x='Percent', y='Dataset',  hue='Type', hue_order=['struct_ii','struct_iii','GC30'],
            data=test, orient='h')
plt.grid(axis='x')

# %% [code]
excluded = ['Chari2015-293T-Cas9','Tálas2021-NEB5-alpha-Cas9','Tálas2021-NEB5-alpha-Cas9-HF1'] 
f, axes = plt.subplots(1,2, figsize=(8,10), sharey=True)

temp = pd.DataFrame(columns=["Dataset_Cas9", "over/under", "Pearson"])

for i, ds in enumerate(mergedLibs[~mergedLibs['Dataset_Cas9'].isin(excluded)].sort_values(['Genome','Year'])['Dataset_Cas9'].unique()):
    tempDF = mergedLibs[mergedLibs['Dataset_Cas9']==ds].copy(deep=True)
    
    temp.loc[temp.shape[0]] = [ds, 'all', pearsonr(tempDF['Activity'], tempDF['MFE_gRNA_G'])[0]]
    temp.loc[temp.shape[0]] = [ds, '> -5', pearsonr(tempDF[tempDF['MFE_gRNA']>-5]['Activity'], tempDF[tempDF['MFE_gRNA']>-5]['MFE_gRNA'])[0]]
    temp.loc[temp.shape[0]] = [ds, '<= -5', pearsonr(tempDF[tempDF['MFE_gRNA']<=-5]['Activity'], tempDF[tempDF['MFE_gRNA']<=-5]['MFE_gRNA'])[0]]

sns.barplot(y='Dataset_Cas9', x='Pearson', hue='over/under', orient='h', ax = axes[0], #color='grey',
            data=temp[temp['over/under']!='all'])

temp = pd.DataFrame(columns=["Dataset_Cas9", "over/under", "Pearson"])

for i, ds in enumerate(mergedLibs[~mergedLibs['Dataset_Cas9'].isin(excluded)].sort_values(['Genome','Year'])['Dataset_Cas9'].unique()):
    tempDF = mergedLibs[mergedLibs['Dataset_Cas9']==ds].copy(deep=True)
    
    temp.loc[temp.shape[0]] = [ds, 'all', pearsonr(tempDF['Activity'], tempDF['Duplex Stability'])[0]]
    try: # small datasets don't have gRNA below the cutoffs
        temp.loc[temp.shape[0]] = [ds, '> -15', pearsonr(tempDF[tempDF['Duplex Stability']>-15]['Activity'], tempDF[tempDF['Duplex Stability']>-15]['Duplex Stability'])[0]]
    except:
        temp.loc[temp.shape[0]] = [ds, '> -15', 0]    
    try: # small datasets don't have gRNA below the cutoffs
        temp.loc[temp.shape[0]] = [ds, '<= -15', pearsonr(tempDF[tempDF['Duplex Stability']<=-15]['Activity'], tempDF[tempDF['Duplex Stability']<=-15]['Duplex Stability'])[0]]
    except:Duplex Stability
        temp.loc[temp.shape[0]] = [ds, '<= -15', 0]    


sns.barplot(y='Dataset_Cas9', x='Pearson', hue='over/under', orient='h', ax = axes[1],
            data=temp[temp['over/under']!='all'])
axes[0].grid(axis='x')
axes[1].grid(axis='x')

# %% [markdown]
# Supplemental Figure 1

# %% [code]
x = mergedLibs['Dataset_Cas9'].value_counts()


datasetList = ['Wang2019-293T', 'Schwartz2019-PO1f','Guo2018-MG1655',
               'Kim2019-SA-lenti293T','Moreb2017-BW25113']


f, axes = plt.subplots(5,3, figsize=(15,20), sharex='col', sharey='row')

cutoff = -5
GCcutoff = 30
duplexCutoff = -15

for i, ds in enumerate(datasetList):
    sns.lineplot(x='GC_0:20', y='Activity', hue='Cas9 Variant', hue_order=['Cas9','Cas9-HF1','eCas9'],
                 data=mergedLibs[mergedLibs['Dataset']==ds], ci='sd', ax=axes[i,0])
    axes[i,0].set_title(ds)
    axes[i,0].legend(loc='lower left')
    axes[i,0].axvline(GCcutoff, ls='--', color='k')
    if i < 4 or i > 4:
        dsCas = ds+'-Cas9'
        count = mergedLibs[(mergedLibs['Dataset_Cas9']==dsCas) & (mergedLibs['GC_0:20']<=GCcutoff)].shape[0]
        totalcount = mergedLibs[(mergedLibs['Dataset_Cas9']==dsCas)].shape[0]
        percent= (count/totalcount)*100
        axes[i,0].text(0.05, 0.85, f"{count} gRNA out of \n{totalcount} ({percent: .1f}%) less than {GCcutoff}",
                    horizontalalignment='left', verticalalignment='center', transform=axes[i,0].transAxes)
    else:
        dsCas = ds+'-Cas9-HF1'
        count = mergedLibs[(mergedLibs['Dataset_Cas9']==dsCas) & (mergedLibs['GC_0:20']<=GCcutoff)].shape[0]
        totalcount = mergedLibs[(mergedLibs['Dataset_Cas9']==dsCas)].shape[0]
        percent= (count/totalcount)*100
        axes[i,0].text(0.05, 0.85, f"{count} gRNA out of \n{totalcount} ({percent: .1f}%) less than {GCcutoff}",
                    horizontalalignment='left', verticalalignment='center', transform=axes[i,0].transAxes)
 
    
    sns.lineplot(x='Duplex Stability', y='Activity', hue='Cas9 Variant', hue_order=['Cas9','Cas9-HF1','eCas9'],
                 data=mergedLibs[mergedLibs['Dataset']==ds], ci='sd', ax=axes[i,1])
    axes[i,1].set_title(ds)
    axes[i,1].legend(loc='lower left')
    axes[i,1].axvline(duplexCutoff, ls='--', c='k')
    
    if i < 4 or i > 4:
        dsCas = ds+'-Cas9'
        count = mergedLibs[(mergedLibs['Dataset_Cas9']==dsCas) & (mergedLibs['Duplex Stability']<=duplexCutoff)].shape[0]
        totalcount = mergedLibs[(mergedLibs['Dataset_Cas9']==dsCas)].shape[0]
        percent= (count/totalcount)*100
        axes[i,1].text(0.05, 0.85, f"{count} gRNA out of \n{totalcount} ({percent: .1f}%) less than {duplexCutoff}",
                    horizontalalignment='left', verticalalignment='center', transform=axes[i,1].transAxes)
    else:
        dsCas = ds+'-Cas9-HF1'
        count = mergedLibs[(mergedLibs['Dataset_Cas9']==dsCas) & (mergedLibs['Duplex Stability']<=duplexCutoff)].shape[0]
        totalcount = mergedLibs[(mergedLibs['Dataset_Cas9']==dsCas)].shape[0]
        percent= (count/totalcount)*100
        axes[i,1].text(0.05, 0.85, f"{count} gRNA out of \n{totalcount} ({percent: .1f}%) less than {duplexCutoff}",
                    horizontalalignment='left', verticalalignment='center', transform=axes[i,1].transAxes)
            
    sns.lineplot(x='Duplex Stability', y='Activity', hue='Cas9 Variant', hue_order=['Cas9','Cas9-HF1','eCas9'],
                 data=mergedLibs[(mergedLibs['Dataset']==ds) & (mergedLibs['GC_0:20']>GCcutoff)], ci='sd', ax=axes[i,2])
    axes[i,2].set_title(ds)
    axes[i,2].legend(loc='lower left') 
    
```
